# Integration of homeostatic and adaptive oxidative responses by a putative co-chaperone, Wos2, drives fungal virulence in cryptococcosis

**DOI:** 10.1101/2023.04.03.535320

**Authors:** Brianna Ball, Arjun Sukumaran, Samanta Pladwig, Samiha Kazi, Norris Chan, Manuela Modrakova, Jennifer Geddes-McAlister

## Abstract

The increasing prevalence of invasive fungal pathogens are dramatically changing the clinical landscape of infectious diseases and are an imminent burden to public health that lack the resources (i.e., robust antifungals) to tackle this threat. Specifically, the human opportunistic pathogen, *Cryptococcus neoformans,* expresses elaborate virulence mechanisms and is equipped with sophisticated adaptation strategies to survive in harsh host environments. In this study, we extensively characterize Wos2, an Hsp90 co-chaperone homologue, featuring bilateral functioning for both cryptococcal adaptation and virulence strategies. Here, we evaluated the proteome and secretome signatures of Wos2 in enriched and infection-mimicking conditions to reveal a Wos2-dependent regulation of oxidative stress response. The *wos2*Δ strain reports defective intracellular and extracellular antioxidant protection systems measurable through a decreased abundance of critical antioxidant enzymes and reduced growth in the presence of peroxide stress. Additional Wos2-associated stress phenotypes were observed upon fungal challenge with heat shock, osmotic, and cell wall stressors. We demonstrate the importance of Wos2 for *C. neoformans* intracellular lifestyle during *in vitro* macrophage infection and provide evidence for *wos2*Δ reduced phagosomal replication levels. Accordingly, *wos2*Δ featured significantly reduced virulence in a murine model of cryptococcosis. Our study highlights a vulnerable point in the fungal chaperone network that offers a powerful druggable opportunity to interfere with both virulence and fitness.

**Author Summary:** The global impact of fungal pathogens, both emerging and emerged, is undeniable and the alarming increase in antifungal resistance rates hampers our ability to protect the global population from deadly infections. For cryptococcal infections, a limited arsenal of antifungals and resistance demands alternative therapeutic strategies, including an anti-virulence approach, which disarms the pathogen of critical virulence factors, empowering the host to remove the pathogen and clear the infection. To this end, we apply state-of-the-art mass spectrometry-based proteomics to interrogate the impact of a recently defined novel co-chaperone, Wos2, towards cryptococcal virulence using *in vitro* and *in vivo* models of infection. We defined global proteome and secretome remodeling driven by the protein and uncovered a novel role in modulating the fungal oxidative stress response. Complementation of the proteome findings with *in vitro* infectivity assays demonstrated a protective role for Wos2 within the macrophage phagosome, influencing fungal replication and survival. These results underscore differential cryptococcal survivability and weakened patterns of dissemination in the absence of *wos2*. Overall, our study establishes Wos2 as an important contributor to fungal pathogenesis and warrants further research into critical proteins within global stress response networks as potential druggable targets to reduce fungal virulence and clear the infection.

## Introduction

*Cryptococcus neoformans* is an opportunistic fungal pathogen responsible for the highly invasive and difficult-to-manage disease, cryptococcosis. This disease currently inflicts over 230,000 individuals annually and results in 19% of AIDS-related mortality (1,2). Invasive human fungal pathogens are equipped with important virulence determinants for survival within the adverse mammalian host environment, challenging pathogens with an elevated body temperature, effective immune system, and nutrient deprivation. Recently, we applied mass spectrometry-based proteomics to investigate the adaption of *C. neoformans* under nutrient limitation of the essential metal, zinc (3). We observed zinc-associated regulation of Wos2, an uncharacterized Hsp90 co-chaperone homologue, and *in vitro* characterization defined a novel role for the protein in virulence factor modulation (i.e., thermotolerance, zinc utilization, melanin pigmentation, capsule production).

*C. neoformans* Wos2 presents substantial homology to the conserved p23 co-chaperone expressed widely across eukaryotes (4). p23 features considerable chaperone-independent functions, including ribosome biogenesis, Golgi operations and DNA repair activities (5), and is a central component of Hsp90 machinery (6). Hsp90 is a well-characterized heat shock protein (HSP) with supported connections to fungal pathogenesis through expansive interaction networks (i.e., interacting with approximately 10% of all yeast proteins) (7,8). For instance, defined interactions between Hsp90 and a co-chaperone, Stg1, orchestrate morphogenesis and drug resistance in the human fungal pathogen, *Candida albicans* (17). In addition, recent studies highlighted the critical role of p23 in stabilizing the complex that Hsp90 forms with client proteins, demonstrating that p23 is essential for general Hsp90 functioning regardless of the respective client protein (9,10). Moreover, in the model fungal organism *Neurospora cassa,* deletion of the Hsp90 co-chaperones, p23, Sti1 and Aha1 resulted in hypersensitivity to azoles and heat (11).

Given the importance of co-chaperones across a spectrum of fungal pathogens and connections to antifungal susceptibility, the characterization of novel co-chaperones suggests an important avenue of discovery to regulate the prevalence and outcome of fungal infections. In this study, we explore the connection across co-chaperones, virulence, and anti-virulence targets using high-resolution quantitative proteomic profiling. We reveal extensive proteome and secretome remodeling during infection-mimicking conditions compared to enriched environments, indicating the requirement for Wos2 to drive intra- and extracellular homeostasis during infection-related stress. We established that Wos2 regulates *C. neoformans* response to oxidative stress and adaptation during prolonged infectious states within macrophages, influencing fungal replication and survival. Furthermore, this is the first report that supports Wos2 as a novel virulence factor that, upon deletion, significantly attenuates a murine model of cryptococcosis, further supporting the protein as a novel target with therapeutic intervention potential.

## Results

### Culture conditions promote cellular proteome remodeling for *wos2*Δ

To gain insight into the global alterations of the co-chaperone Wos2 on *C. neoformans* proteome, we compared protein production profiles between *C. neoformans* wild-type (WT) and a *wos2* deletion strain. We investigated the connection between Wos2 and fungal adaptation to a host-like environment by profiling the protein-level changes within the cellular proteome (cell pellet) of cells grown to mid-log phase under enriched (i.e., YPD) and infection-mimicking (i.e., low-iron media, LIM) conditions (Fig 1) (12,13).

**Fig 1.**
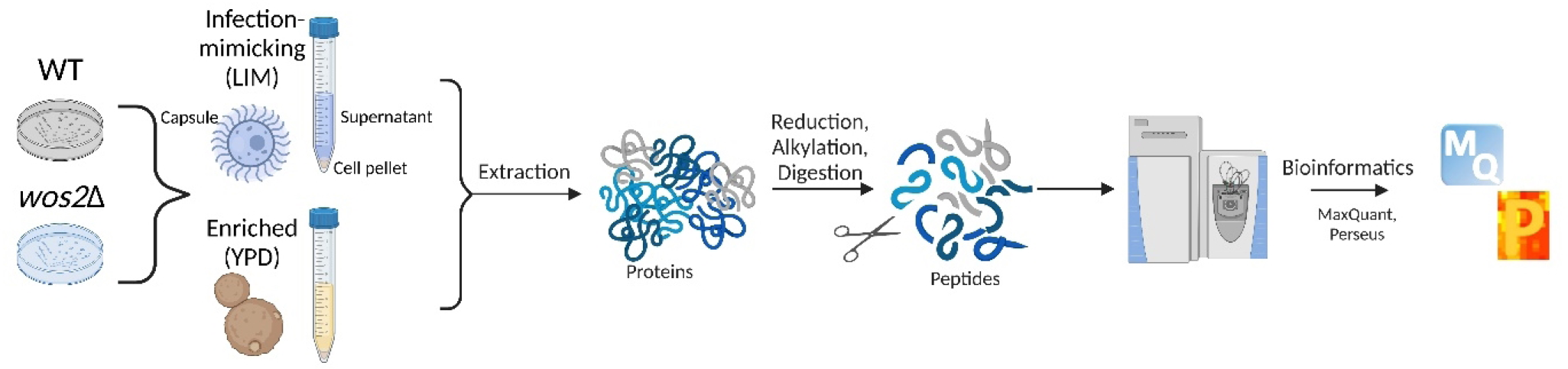
Workflow for mass spectrometry-based proteomics profiling. *C. neoformans* WT and *wos2*Δ strains were cultured in infection-mimicking (low-iron media, LIM) or enriched (yeast peptone dextrose, YPD) media and subjected to our total and secretome protein extraction protocol (e.g., sonication with detergent), followed by protein reduction, alkylation, and digestion, and measurement on the mass spectrometer. Data analysis, visualization, and statistical processing performed using MaxQuant and Perseus (55,58). The experiment was completed in biological quadruplicate. Figure generated with Biorender.com.

Our analysis detected 3420 unique proteins (2933 proteins after valid value filtering) across the samples, representing approximately 46% of *C. neoformans* proteome. Of these, the YPD-enriched proteome consisted of 3065 proteins shared between WT and *wos2*Δ, whereas 80 proteins were detected only in the WT proteome and 67 proteins identified solely within *wos2*Δ (Fig 2A). Conversely, the infection-mimicking proteome consisted of 1434 proteins identified from both strains, whereas the WT and *wos2*Δ proteomes identified 113 and 607 unique proteins, respectively, with 1434 core proteins shared between the strains (Fig 2B). Next, we classified the unique proteins based on Gene Ontology of Biological Processes (GOBP) and observed a common abundance of proteins associated with biosynthetic and metabolic processes as well as uncharacterized roles with detection of transport-associated proteins unique to WT. Similarly, infection-mimicking conditions revealed consistent representation of protein categories between the strains, including an anticipated increase in transport-associated proteins compared to enriched conditions given the need for nutrient acquisition from the environment for both strains. Additionally, an emphasized response to cellular assembly and growth for WT was defined compared to the *wos2* deletion strain.

**Fig 2.**
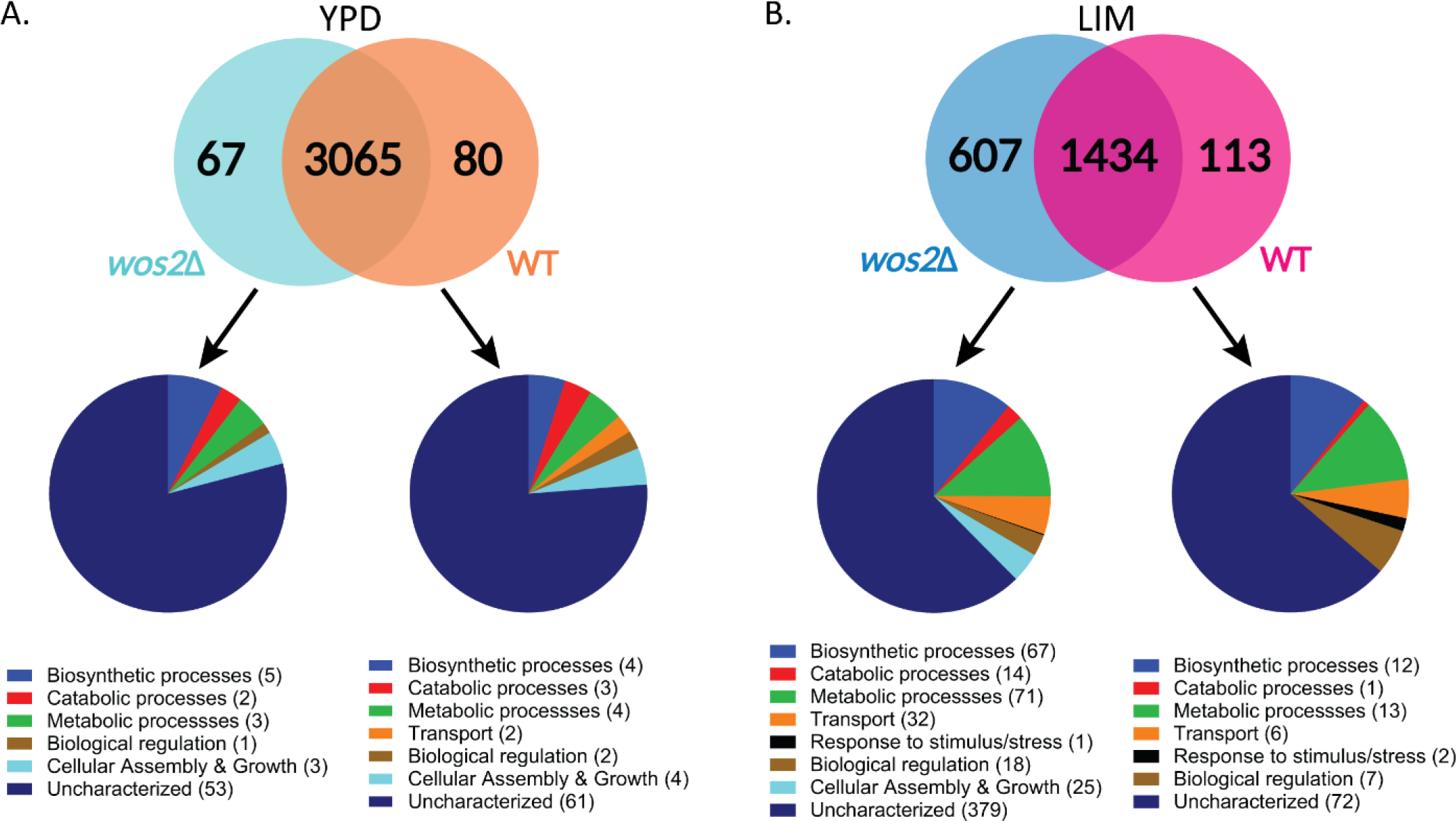
Wos2-dependent cellular proteome signatures between enriched and infection-mimicking environments. A) Venn diagram for number of unique proteins identified in the YPD (enriched) cellular proteome between *C. neoformans* WT (orange; 80) and *wos2*Δ (turquoise; 67) strains with 3065 proteins commonly identified. The distribution of Gene Ontology Biological Processes (GOBP) terms for identified unique fungal proteins are shown exclusive to each strain. B) Venn diagram for number of unique proteins identified in the LIM (infection-mimicking) cellular proteome between *C. neoformans* WT (pink; 113) and *wos2*Δ (blue; 607) strains with 1434 proteins commonly identified. Experiment performed in biological quadruplicate. YPD: yeast peptone dextrose; LIM: low-iron media.

A principal component analysis (PCA) of the cellular proteome indicated the largest component of separation as fungal growth conditions (component 1, 69.5%), with the second component of separation attributed to the absence of *wos2* (component 2, 7.1%) (Fig 3A). A comparison of significant differences in protein abundance between WT and *wos2*Δ under enriched conditions defined eight proteins significantly more abundant in WT and two proteins with significantly higher abundance in the mutant (Fig 3B; S1 Table). Importantly, Wos2 (CNAG_07558) was > 7.5-fold (log_2_) in the WT strain compared to the mutant strain, confirming disruption. We also observed an increase in abundance of proteins within the WT strain important for reactive oxygen species (ROS) detoxification, including the well-characterized antioxidant defense protein, catalase 3 (CAT3, CNAG_00575; > 2.4-fold [log_2_]), as well as an uncharacterized membrane protein (CNAG_03394; >4.5-fold [log_2_]) with predicted peroxisomal localization, an organelle essential for sequestering diverse oxidative reactions (14–16). Furthermore, two uncharacterized proteins (CNAG_04585; >2.1-fold [log_2_], CNAG_01387; >2.3-fold [log_2_]) were more abundant in the WT strain along with proteins involved in RNA processing, including nuclease I (CNAG_00264; > 2.2-fold [log_2_]), a ribosome assembly factor (midasin, CNAG_00848; >2.3-fold [log_2_]), and She3 (SWI5-dependent HO expression protein 3, CNAG_03966; >2.7-fold [log_2_]) involved in mRNA localization (17,18). Conversely, in the *wos2*Δ strain, two uncharacterized membrane-bound proteins were more abundant, including an uncharacterized protein involved in transport to the plasma membrane (CNAG_07020; >2.1-fold [log_2_]) and a protein with predicted phospholipid biosynthesis activity (CNAG_04522; >2.9-fold [log_2_]), suggesting alterations in the cellular organization of both phospholipid membranes and their proteinaceous components in the absence of Wos2.

**Fig 3.**
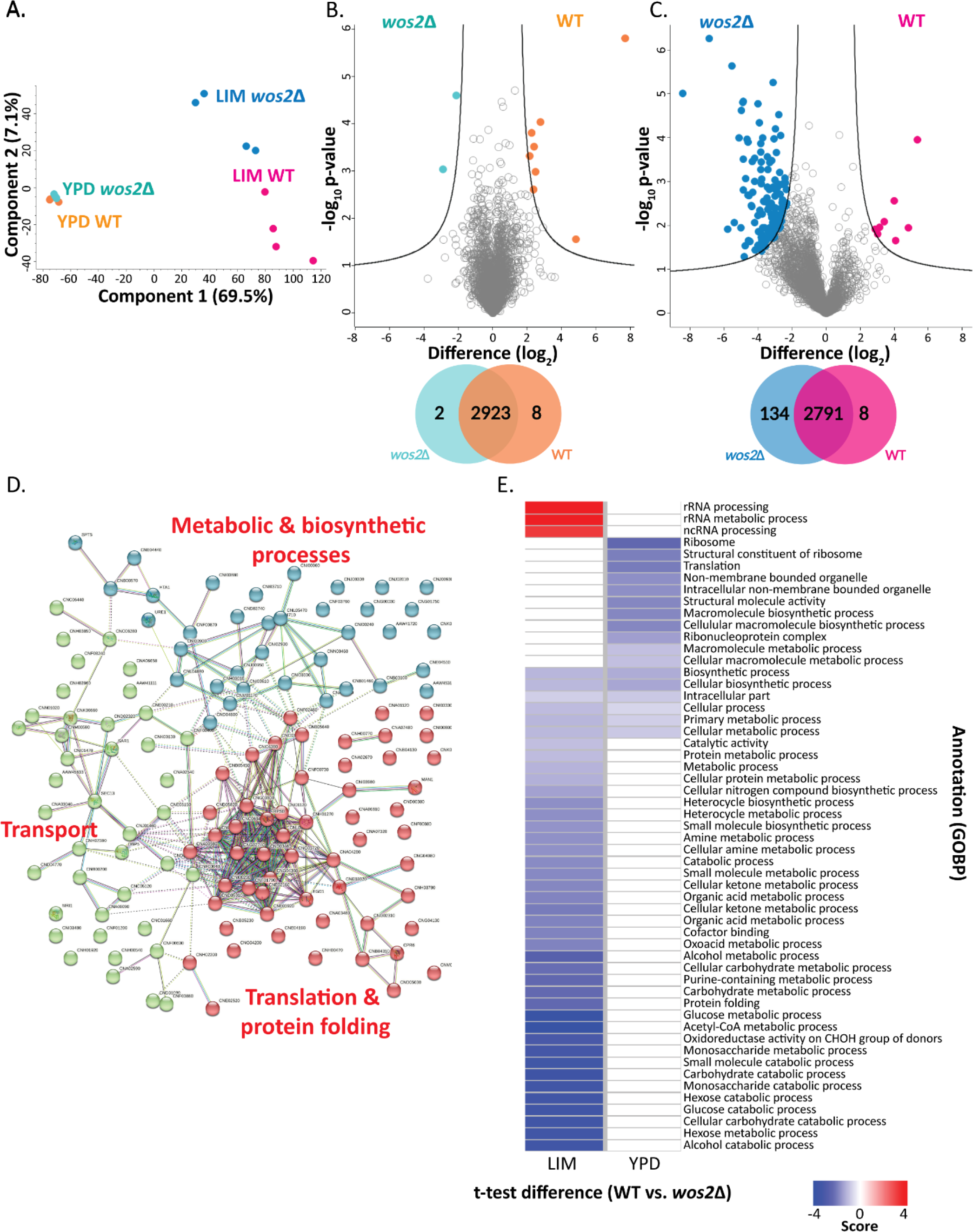
Wos2 remodeling of the global fungal proteome. A) Principal component analysis for YPD (enriched) and LIM (infection-mimicking) conditions. B) Volcano plot comparing proteins identified under enriched (i.e., YPD) fungal growth between WT and *wos2*Δ. Highlighted proteins depict significant changes in protein abundance upon strain comparison; Venn diagram depicts the number of significantly different proteins identified in each condition. C) Volcano plot comparing proteins identified under infection-mimicking (i.e., LIM) conditions between WT and *wos2*Δ. Highlighted proteins depict significant changes in protein abundance upon strain comparison; Venn diagram depicts the number of significantly different proteins identified in each condition. Students *t* test, *P* < 0.05; FDR = 0.05; S_0_ = 1. D) STRING analysis for fungal proteins with significant increase in abundance within *wos2*Δ under infection-mimicking conditions (LIM). Cluster themes were classified based on gene names and protein function annotations within STRING. E) 1D annotation enrichment for proteins defined within categories based on Gene Ontology Biological Processes (GOBP). Students *t* test, *P* < 0.05; FDR = 0.05. Experiment performed in biological quadruplicate.

Given the stability of the total proteome under enriched conditions but the defined role of Wos2 in fungal virulence factor production (3), we next assessed the proteome remodeling under infection-mimicking conditions. Here, we observed a significant increase in production of eight proteins in the WT strain and a vast difference in the mutant strain, with 134 proteins showing significantly higher production (Fig 3C; S1 Table). Across both enriched and infection-mimicking conditions, the AAA ATPase midasin (CNAG_00848; >2.3-fold [log_2_]) was consistently abundant within the WT strain, with known roles in ribosome assembly, as well as predicted nuclear chaperone duties (17,19). Additionally, of the proteins with increased production in WT, multiple proteins feature a wide range of functions within cellular respiration, including an NADH-ubiquinone oxidoreductase subunit (CNAG_05267; >3-fold [log_2_]) of the electron transport chain (ETC), a mitochondrial pyruvate carrier (CNAG_00092; >3.3-fold [log_2_]), and Oxa1 (CNAG_02769; >4.8 fold [log_2_]), which mediates the assembly of components within the ETC (20). Conversely, a high-level overview of *wos2*Δ significantly different proteins defined clustering into three major categories via STRING analysis (21): translation & protein folding, transport, and metabolic & biosynthetic processes (Fig 3D). Interestingly, the well-defined cytokine-inducing glycoprotein (Cig1), which is an established *C. neoformans* virulence-associated hemophore, important for iron acquisition and production in iron-starved cells (22), was the most abundant protein detected for *wos2*Δ (>8.4-fold [log_2_]), supporting further connection between Wos2 and nutrient limitation and fungal virulence.

Given the diverse impact of Wos2 on proteome regulation under altered media conditions, we performed a 1D annotation enrichment (23) to comprehensively characterize the influence of Wos2 at the categorical level based on GOBP. Under YPD conditions, we observed enrichment of proteins within *wos2*Δ associated with the ribosome, translation, structural activity, and biosynthetic and metabolic processes (Fig 3E); no enrichment of categories for the WT were observed. Under infection-mimicking (LIM) conditions, we observed broader enrichment across protein categories, including rRNA processing and metabolic process, and ncRNA processing for WT. Conversely, for *wos2*Δ, we observed enrichment across 41 categories with an emphasis on molecular metabolic (i.e., hexose, monosaccharide, acetyl-CoA, glucose, etc.) and catabolic (i.e., alcohol, carbohydrate, glucose, etc.) processes, protein folding, and oxidoreductase activity. Together, these data support Wos2 as a central player in fungal adaptation from homeostasis towards stress- and virulence-induced conditions and define remodeling at the protein level to drive these adaptations.

### Wos2 regulates fungal adaptation to heat shock, cell wall, and osmotic stress and drives tolerance to oxidative stress

Fungal pathogens require robust adaptation strategies to endure stressors within hostile environments (e.g., during infection of a human host), and a lack of homeostatic response can lead to cellular dysregulation in intracellular protein transport, disruption of cellular organization, and lethal proteotoxicity (8,24). Given the homology of Wos2 to an Hsp90 co-chaperone and its important roles in proteomic reprogramming during both enriched and infection-mimicking conditions, we aimed to establish the requirement of Wos2 for fungal stress response. Fungal growth on YPD at 37°C was assessed as a control and showed no difference among WT, *wos2*Δ and *wos2*Δ::WOS2 (Fig 4A). For thermal stress, we assessed the heat shock (42°C) sensitivity of *wos2*Δ and revealed a subtle decrease in fungal growth on YPD at elevated temperatures (Fig 4B). Next, we assessed the sensitivity of the *wos2*Δ mutant to osmotic and cell wall stressors. For example, when strains were incubated in the presence of caffeine, an effector of signal transduction and cell wall integrity (25), no divergences in growth were observed (Fig. 4C). In contrast, *wos2*Δ featured a noticeable growth inhibition compared to WT when grown in the presence of the cell membrane stressor, SDS (Fig. 4C). Fungal growth on multiple osmotic stressors, including NaCl and KCl, revealed a more pronounced growth sensitivity and a subtle growth impairment upon exposure compared to WT (Fig 4D), suggesting that Wos2 is important for diverse stress responses.

**Fig 4.**
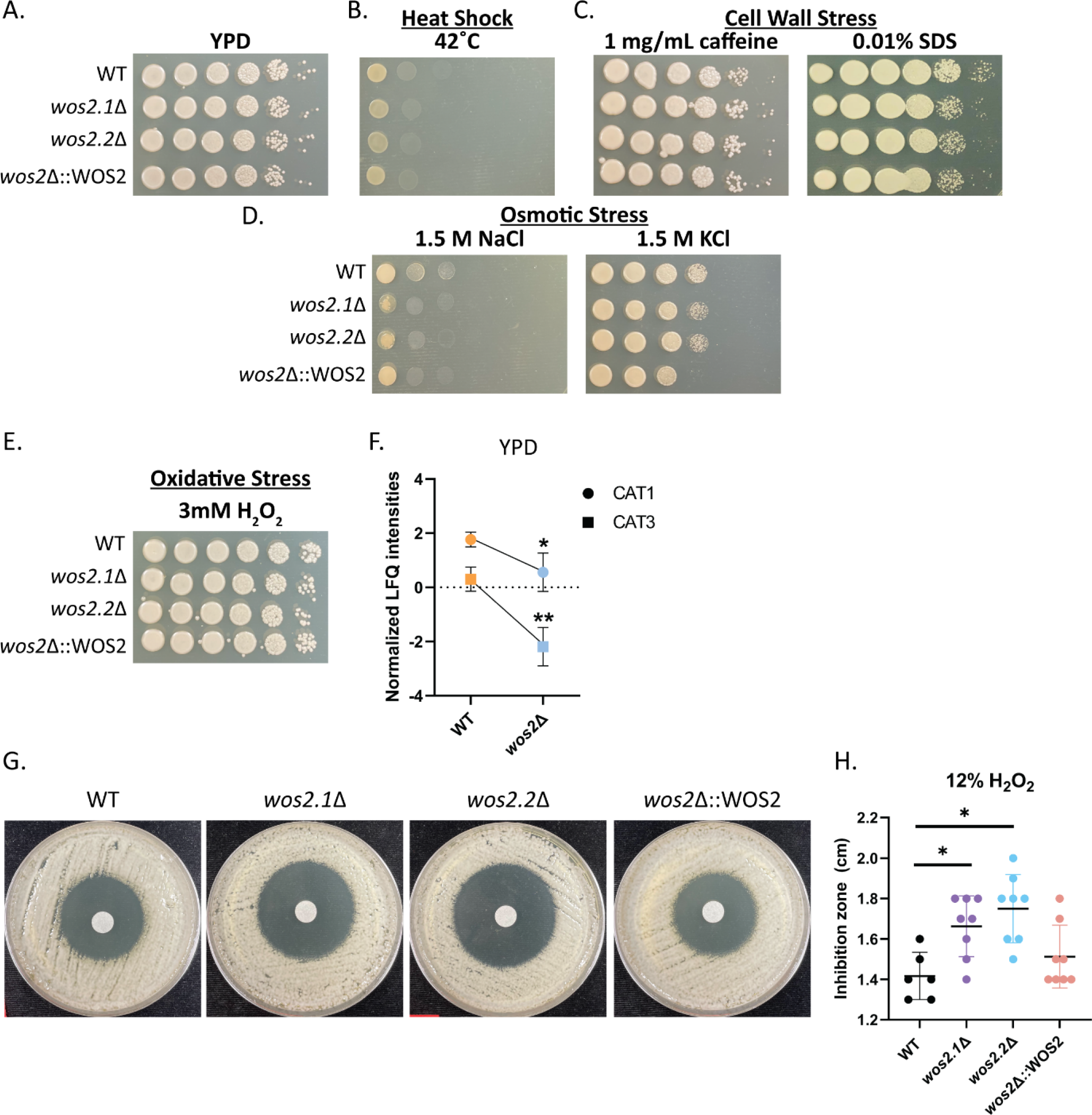
Wos2 influences fungal susceptibility to multiple stressors. A-D) Phenotypic screening of *wos2*Δ mutant under A) rich conditions, and against B) heat (i.e., 42°C), C) cell wall (i.e., 1 mg/mL caffeine, 0.01% SDS), and D) osmotic (i.e., 1.5 M NaCl, 1.5 M KCl) stressors. Serial dilutions of strains were spotted onto YPD and YPD supplemented with stressor and incubated at 37°C for 48 h, unless otherwise stated. E) Phenotypic screening of *wos2*Δ mutant against oxidative (i.e., 3 mM H_2_O_2_) stress. Dilution plate experiments completed in biological triplicate and technical duplicate. F) Normalized mean label free quantitative (LFQ) intensities for *C. neoformans* CAT1 and CAT3 proteins identified by proteomic profiling of YPD-enriched conditions. Experiment completed in biological quadruplicate. G) H_2_O_2_ disc diffusion across WT, *wos2*Δ, and *wos2*Δ::WOS2 strains in the presence of 12% H_2_O_2_. Experiment performed in biological quadruplicate and technical duplicate. H) Catalase production assessed from H_2_O_2_ disc diffusion assay and measured based on zone of inhibitions. Statistical analysis using Students *t* test: *, *P* < 0.05; **, *P* ≤ 0.001.

Given our consistent observations of stress response regulation across the Wos2-dependent proteomes, including proteins associated with oxidative stress (i.e., catalases) and a peroxisomal membrane protein, we evaluated the impact of ROS on the *wos2*Δ strain. As anticipated, growth of the fungal strains in the presence of 3 mM H_2_O_2_ (an oxidative stressor) showed a minor reduction in growth for *wos2*Δ vs. WT (Fig 4E). These data are supported by proteomics profiling defining a significant reduction in catalase (CAT1, CAT3) production in the *wos2*Δ vs. WT strains (Fig 4F), sustaining increased ROS protection for the WT strain. To further elucidate these findings, we evaluated catalase production in the presence of oxidative stress (e.g., H_2_O_2_) and revealed a difference in catalase production between the WT and complement strains compared to *wos2*Δ observed by a zone of inhibition (Fig 4G). A quantitative assessment of the zone of inhibition measurements revealed a significant increase in protective catalase production for the WT strain as evident by a smaller zone of clearance compared to the *wos2*Δ strains (Fig 4H). Collectively, these results support Wos2 as an important mediator of fungal adaptation to ROS.

### Secretome profiling further elaborates on the role of Wos2 in oxidative stress response

The extracellular environment serves as an important interaction point for fungal pathogens to respond to the milieu of host-generated ROS as a first line of defense and subsequent adaptation (26). Given the pronounced Wos2-signature within the infection-mimicking cellular proteome, we explored if Wos2 exhibits similar control over the extracellular environment in host-like conditions. Here, we profiled the secretome across WT and *wos2*Δ strains when grown in LIM. A PCA plot indicated the largest component of separation was under genetic regulation (i.e., deletion of *wos2*) (component 1, 56.4%) with a second component of distinction defined by biological variability (component 2, 18.5%) (Fig 5A). A volcano plot comparison revealed distinct Wos2-signatures of the fungal extracellular environment under infection-mimicking conditions with five proteins significantly more abundant in WT vs. two proteins with significantly higher abundance in the deletion strain (Fig 5B; S2 Table). For WT, we observed a significant increase in abundance of the well-characterized *C. neoformans* virulence factor, superoxide dismutase (SOD, CNAG_01019; > 1.3-fold [log_2_]), a key protein for detoxifying oxygen radicals previously reported to be packaged cargo within fungal extracellular vesicles (27,28). These data solidify the role of Wos2 in universal oxidative stress response. We also identified β-1,3-glucanase (CNAG_02860; >2.9-fold [log_2_]), a major structural constituent of the fungal cell wall dependent on functional glucanases for cell wall biogenesis (29), corroborating the observed loss of cell wall integrity induced by membrane stress within the *wos2*Δ (i.e., SDS, Fig 4C). In addition, an uncharacterized protein (CNAG_02843; > 1.0-fold [log_2_]), a nucleosome assembly protein (CNAG_02091; > 1.2-fold [log_2_]) and elongation factor 1-gamma (CNAG_00417; > 2.2-fold [log_2_]) were significantly more abundant in WT. Conversely, we identified a voltage-gated potassium channel subunit (CNAG_04209; > 2.0-fold [log_2_]) with predicted importance in regulating membrane potential (30), as well as a previously reported moonlighting protein, transaldolase (CNAG_01984; > 1.4-fold [log_2_]) (31), to be more abundant in the *wos2*Δ strain. Altogether, these results support a role for Wos2 in modulating the secretory profile of *C. neoformans* under nutrient limitation for preparation against host defenses, including ROS.

**Fig 5.**
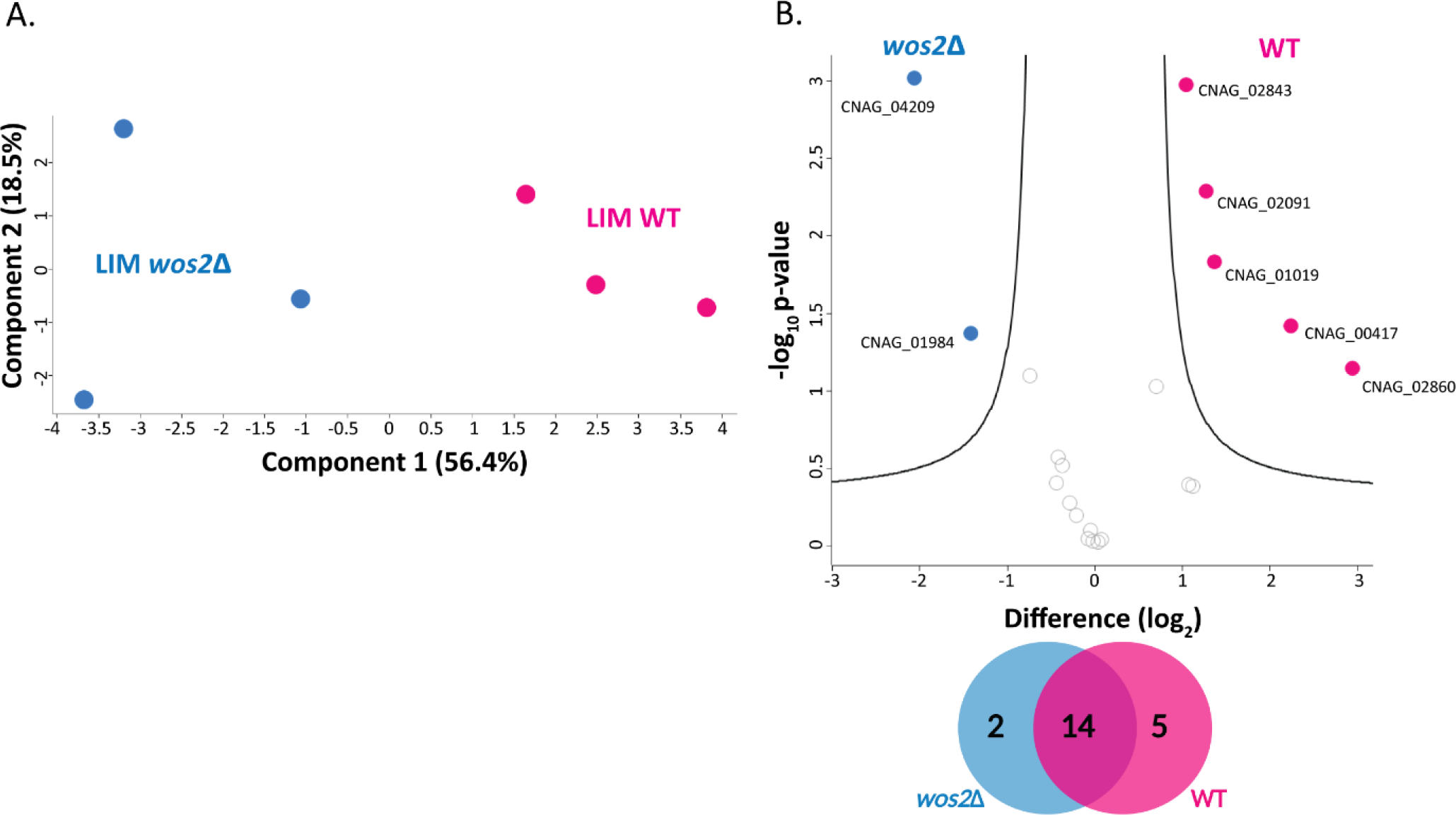
Secretome profiling of Wos2 in infection-mimicking conditions. A) Principal component analysis of experiment profiled in LIM (infection-mimicking) conditions between WT and *wos2*Δ strains. B) Volcano plot comparing all proteins identified under infection-mimicking (i.e., LIM) conditions between WT and *wos2*Δ. Highlighted proteins depict significant changes in protein abundance upon strain comparison; Venn diagram depicts the number of significantly different proteins identified in each condition. Students *t* test, *P* < 0.05; FDR = 0.05; S_0_ = 1. Experiment performed in biological triplicate.

### Wos2 mediates intracellular fungal survival and replication within macrophages and attenuates fungal virulence

Given the role of Wos2 in modulating cryptococcal response to ROS, we assayed the localization during macrophage infection to reveal potential infection-associated functioning. Through immunomicroscopy of a Wos2-FLAG tagged strain, we observed Wos2 production in the presence of macrophages within the fungal cell cytoplasm (Fig. 6A). These data align with defined localization for other co-chaperones, such as Hsp90, which is localized cytosolically and to the cell wall (32).

**Fig 6.**
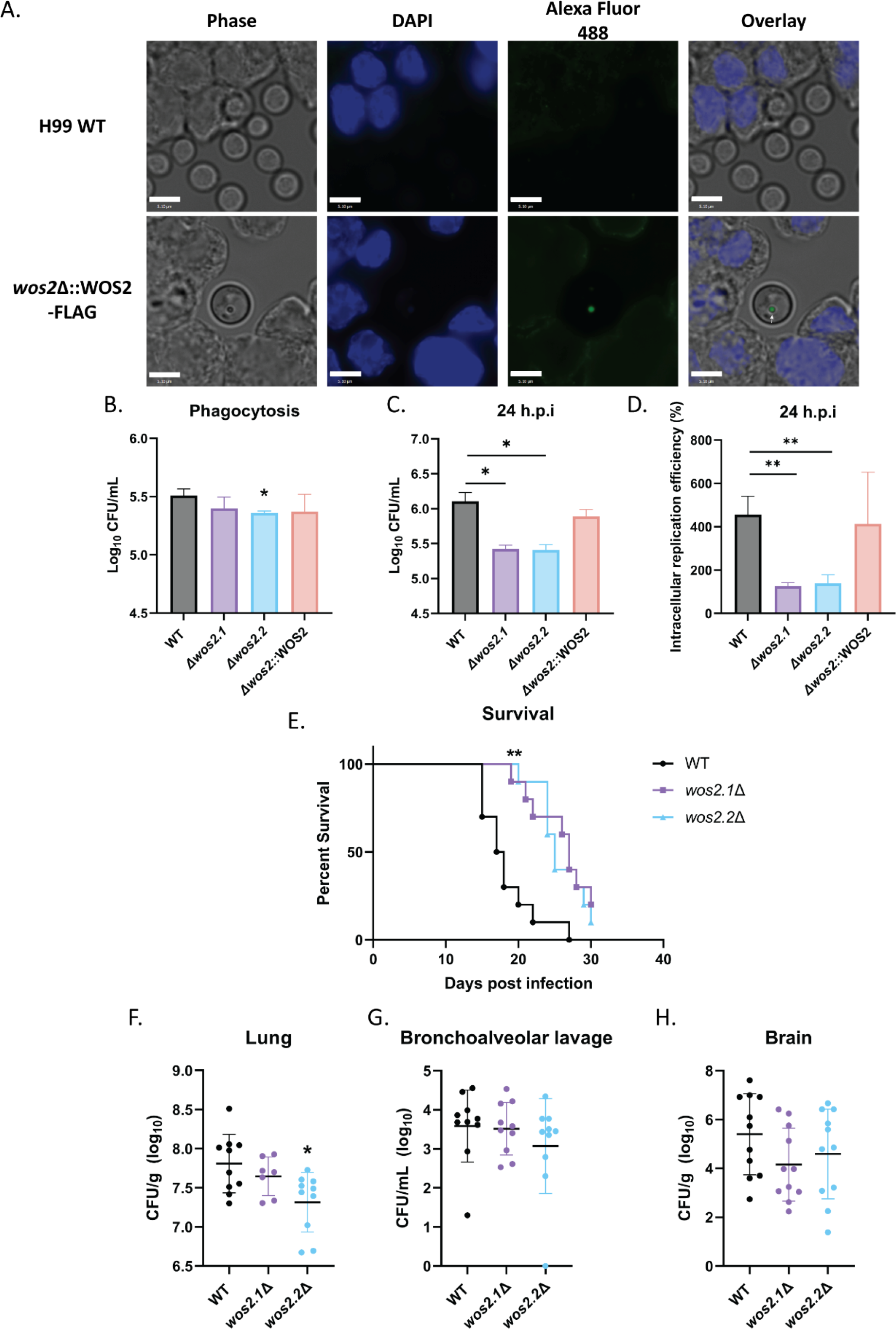
Characterization of Wos2 during *in vitro* and *in vivo* infection assays. A) Fluorescence microscopy of *C. neoformans wos2*Δ::WOS2-FLAG co-cultured with macrophage (MOI 100:1). Images captured at 3 h.p.i with DAPI and Alexa Fluor 488 (FLAG). Scale bar 5.1 µM. B) Phagocytosed fungal cells were quantified by co-culturing macrophages with WT and *wos2*Δ strains for 3 h. Infected cells were washed six times with PBS, lysed and plated for CFUs. C) Fungal burden was assessed following initial 3 h co-culture and six PBS washes to remove extracellular and non-adhered fungal cells, followed by maintenance of infected macrophages in DMEM supplemented with 20 μg/mL fluconazole. At 24 h.p.i, host cells were lysed and plated for CFUs. D) Intracellular replication efficiencies were calculated by quantifying the number of CFUs in the supernatant and host cell at 24 h.p.i. Statistical analysis using Students *t* test: *, *P* < 0.01; **, *P* ≤ 0.005. Experiments completed in biological triplicate and technical duplicate. E) BALB/c mice infected with *C. neoformans* WT and *wos2*Δ succumbed to infection or *wos2*Δ mutant survived to assay endpoint (i.e., 30 days). Differences in survival statistically tested using a log-rank (Mantel-Cox) test (**, *P* ≤ 0.001). F-H) Fungal burden from lung (F), bronchoalveolar lavage fluid (G) and brain (H) determined measuring CFUs. Statistical analysis using Students *t-*test (*, *P* < 0.01).

Next, given our defined role for Wos2 in oxidative stress and fungal virulence, and the ability of *C. neoformans* to interact and survive within macrophages, especially within the harsh environment of the phagosome (e.g., ROS, reactive nitrogen species, nutrient starvation and acidic pH) (26), we speculated that *wos2*Δ strains would be compromised in ROS detoxification abilities and demonstrate increased sensitivity within the phagosome environment. To test our hypothesis, we first assessed the phagocytic ability of macrophages to engulf *wos2*Δ via co-culturing of the fungal strain with immortalized BALB/c macrophages. We observed a slight reduction in phagocytosis for the *wos2*Δ strains (significant reduction for one independent mutant), suggesting Wos2 may influence macrophage engulfment but likely not to biologically relevant levels (Fig 6B). Notably, our previous observations of enlarged *wos2*Δ cell size may support such impeded fungal uptake by macrophages (3). We then assessed the importance of Wos2 within a prolonged infectious state by quantifying intracellular populations of *C. neoformans* upon macrophage exposure. At 24 h.p.i, we observed a significant reduction in fungal burden relative to WT for the *wos2*Δ strains (Fig 6C). To address if these variances in fungal survival were associated with differing macrophage phagocytic uptake or intracellular replication, we applied a fluconazole protection assay at 24 h.p.i. (33). Assessment of CFU counts from the supernatant and lysed macrophages at 24 h.p.i. normalized to CFUs of the phagocytosed fungal strain identified a significant impairment in replication efficiencies for *wos2*Δ compared to WT (Fig 6D). To verify that *wos2Δ* viability was not disproportionately reduced relative to WT due to fluconazole susceptibility, growth in DMEM with and without fluconazole showed no significant differences in OD_600nm_ measurements (final average measurement of 0.15 for WT and *wos2*Δ strains). Critically, these data align with our *in vitro* phenotypic assay findings and proteomic results to support a deficit for *wos2*Δ to replicate under ROS-inducing conditions.

Given the *wos2*Δ virulence defect in oxidative stress and impaired affinity for the intracellular macrophage environment, we predicted that *wos2*Δ would display reduced virulence in an inhalation model for cryptococcosis. Following infection, we observed a significant increase in murine survival with the *wos2*Δ strain compared to WT (Fig 6E). Moreover, to assess if differences in survival were attributed to reduced fungal burden, we quantified a significant decrease of fungal cells within lungs infected with one independent mutant (*wos2.2*Δ) (Fig 6F). Notably, the second independent mutant (*wos2.1Δ*) showed the same trend in reduced fungal burden but not to a significant level. Bronchoalveolar lavage did not reveal any significant changes in fungal burden compared to WT (Fig 6G), as also noted within the brain (Fig 6H), although a reduction in fungal counts for both organs was detectable. Taken together, these data support Wos2 as an important contributor to *C. neoformans* virulence with an impaired ability to cause disease in an *in vivo* model of cryptococcosis without concretely impacting the fungal burden across the lungs and brain.

## Discussion

In this study, we apply high-resolution mass spectrometry-based proteomics to investigate global proteome changes of Wos2, a predicted Hsp90 co-chaperone based on homology, across nutrient-rich and infection-mimicking conditions. Our approach profiles the involvement of Wos2 in fungal homeostasis and adaptation upon exposure to nutrient limitation and *C. neoformans-*specific virulence-inducing stressors. Across conditions, we reveal a Wos2-dependent protein signature centered on remodeling the proteome and secretome towards homeostatic maintenance featuring an enrichment of proteins involved in defense mechanisms against oxidative stress. Furthermore, we support the detected protein reorganization in *wos2*Δ with phenotypic assays validating involvement in response to thermal, osmotic, oxidative and cell wall stressors. These phenotypic divergences were corroborated within *in vitro* macrophage and *in vivo* murine models, where we observed an impairment of *wos2*Δ for the phagosomal lifestyle, which translated to attenuated fungal virulence. Altogether, we reveal the impressive integration of Wos2 within the fungal stress and adaptation protein network and establish the requirement of Wos2 for fungal virulence.

The role of co-chaperones in fungal virulence is well supported by HSP characterization as primary responders to environmental stress (32,34–36). The relationship between HSPs and fungal virulence is well represented by the Hsp70s and Hsp90s (8). For example, *C. neoformans* depend on Hsp90 machinery for thermotolerance, and Hsp70 is involved in melanin pigmentation (37,38). Additionally, proteomic analysis revealed the packaging of both Hsp70 and Hsp90 into extracellular vesicles along with a myriad of other pathogenesis-related molecules resulting in the construction of ‘virulence bags’ capable of delivering pathogenic cargo across the body during infection (28,32,35). In comparison, Hsp90 of *C. albicans* is vital in commanding yeast to hyphal transitions and regulating drug resistance (39–41). However, HSPs do not function independently; they rely on a complex network of proteins to facilitate client protein folding and modifications. For instance, co-chaperones in the Hsp90 system feature polarizing control over Hsp90-mediated client folding (9). In this study, loss of Wos2 impacted protein folding under LIM conditions, thus classifying Wos2 as a co-chaperone with a distinct stress response signature towards infection-mimicking stress.

A promising approach to combatting the limited antifungal supply is generating novel therapeutics capable of interrupting core fungal stress responses, such as the Hsp90 chaperone. However, due to close evolutionary history, a significant hurdle is overcoming host toxicity (36,42,43). Therefore, elucidating the roles of Wos2 and other co-chaperones offers an untapped resource for drug development and understanding foundational stress responses in fungal pathogens. For instance, hypersensitivity to the Hsp90 inhibitors, geldanamycin and radicicol, was observed in yeast lacking the Wos2 homolog, p23 (44). Furthermore, Hsp90 co-chaperones may feature stress-specific contributions depending on the type of stressor; for instance, p23 deletion in *N. cassa* resulted in hypersensitivity to azoles and heat (11). The major response of HSPs and co-chaperones to mitigating thermal stress is ensuring proper protein folding and protein conformation (8). Therefore, the observed subtle *wos2* specific heat shock sensitivity is characteristic of an impaired Hsp90 co-chaperone network and complements our observed enrichment of GOBP terms associated with translation and protein folding upon *wos2* deletion. Another key chaperone trait in fungal pathogens is the ability to influence azole susceptibility. Specifically, azole-induced stress results from the accumulation of toxic sterol intermediates that compromise cell membrane integrity and result in membrane stress (34,40). Given that Hsp90 mediates fluconazole efficacy in *C. neoformans* (34), our observation of Wos2-required maintenance of membrane integrity under a membrane stressor motivates further investigation into the connection of Wos2 and azole susceptibility. Additionally, we did not distinguish between chaperone-dependent and Wos2-independent functioning; therefore, it would be of future interest to tease apart this protein-protein interaction to elucidate Wos2-centred functions concerning fungal adaption and virulence.

The ability to cause and maintain infection within a host depends on the capacity to endure the released extracellular ROS milieu and the oxidative burst within host immune cells. High levels of oxidative stressors result in catastrophic disruptions in homeostatic cellular functions via the oxidation of proteins, lipids, and nucleic acids (45). Thus, fungal pathogens evolved antioxidant defense mechanisms and chaperoning networks to repair and prevent cellular damage, ensuing multiple safety precautions to maintain fungal survival (46). We revealed that deletion of *wos2* resulted in a defective antioxidant protection system with decreased responsiveness in major antioxidant enzymes (i.e., Sod1, Cat1, Cat3) and a measurable oxidative sensitivity phenotype (i.e., H_2_O_2_). Typically, *C. neoformans* intracellular survival within macrophages is managed by modifying the phagosome to be more hospitable, including the expression of powerful antioxidant strategies (i.e., capsule, melanin, enzymes) to absorb the oxidative burst and promote intracellular replication (27,47,48). Thus, an impaired fungal defense response leads to oxidative susceptibility. For instance, the deletion of *sod1* in *C. neoformans* results in reduced growth rates within macrophages (27). Similarly, Wos2 deletion resulted in decreased fungal burden and intracellular replication levels, suggesting Wos2-dependent oxidative protection promotes an affinity for the phagosomal environment and a satisfactory replication niche.

Furthermore, *C. neoformans* requires immediate protection against host oxidative stress upon initial lung infection due to the high exposure to alveolar macrophages; thus, an adequate anti-oxidative response promotes lung colonization and positive macrophage association supporting extrapulmonary dissemination (49). *C. neoformans* interaction with macrophages contributes to disease persistence by undetected fungal dissemination within phagosomes and fungal-mediated cytotoxicity, including lytic exocytosis from unchecked intracellular replication, leading to the release of fungal cells and death of the host cell (26,50,51). We observed reduced virulence in a mouse model of cryptococcosis with the *wos2*Δ strain compared to WT, with both strains exhibiting similar dissemination profiles based on fungal burden analysis in key organs. We propose that *wos2*Δ strains can transit within macrophages at minor levels without inflicting host-lytic damage due to reduced intracellular replicative abilities but maintain a complete and detectable dissemination profile comparable to WT. Thus, Wos2 is an important component of fungal virulence; however, it is not a requirement to cause disease. This fungal durability may be explained by *C. neoformans* redundant and functionally overlapping assortment of virulence and protection mechanisms. Overall, this highlights the delicate host-fungal interaction balance and how debilitating a significant arm of the fungal stress response network leads to compensatory proteome regulation to maintain control and infection within the host.

## Conclusions

Chaperones are a critical line of fungal protection against cellular proteotoxic damage induced by high temperatures and environmental stressors. This global stress response comprises of overlapping chaperone and co-chaperone networks consisting of many clinically relevant proteins important for multiple facets of fungal disease. Partitioning putative co-chaperones, such as Wos2, as a potential therapeutic avenue facilitates the inhibition of a layer of Hsp90 regulation and the entire co-chaperone stress response system. Therefore, it is important to detail the involvement of a co-chaperone in both fungal virulence and fitness, as the ability of a fungal pathogen to cause disease is fitness conditional. Our study provides a detailed investigation into the co-chaperone Wos2 and defines distinct Wos2-controlled fitness and virulence attributes vital for response to environmental threats and establishing fungal infection.

## Materials and Methods

### Fungal strains, growth conditions and media

*Cryptococcus neoformans* var. *grubii* wild-type (WT) strain H99 (serotype A) was used for all analyses and as a reference strain for mutant construction. *C. neoformans* mutant (*wos2*Δ) and complement strains (*wos2*Δ::WOS2) were constructed using biolistic transformation of constructs amplified using double-joint PCR or Gibson Assembly (NEB), as previously described (primers, strains, and plasmids provided) (3). Two independent mutant strains were constructed for *wos2* (strains listed in S3). Wild-type strain was maintained on yeast peptone dextrose (YPD) agar plates (2% dextrose, 2% peptone, 1% yeast extract, and 1% agar), and mutant and complement strains were maintained on YPD supplemented with 100 μg/mL nourseothricin (NAT) and 200 μg/mL hygromycin B, respectively, at 30°C unless otherwise stated.

For YPD (i.e., enriched) proteomic sample collection, *C. neoformans* strains were inoculated in YPD media overnight at 37°C, followed by subculture into fresh YPD and grown to mid-log phase (approx. 13 h). For infection-mimicking proteomic sample collection, fungal strains were inoculated in YPD media overnight at 37°C, followed by subculture in yeast nitrogen base (YNB) medium with amino acids (BD Difco, Franklin Lakes, NJ) supplemented with 0.05% dextrose overnight. Samples were collected, washed in low iron capsule inducing media (CIM) and sub-cultured in CIM to mid-log phase (approx. 37 h) (12). Proteomic samples were collected in biological quadruplicate.

### Proteomics sample preparation

Samples for mass spectrometry were prepared as previously described (52). Briefly, samples were collected and washed twice in 1x phosphate buffered saline (PBS) and resuspended in 100 mM Tris-HCl (pH 8.5) containing a protease inhibitor cocktail tablet (Roche). Following the addition of sodium dodecyl sulfate (SDS, 2% final concentration), samples were lysed using a probe sonicator (Thermo Fisher Scientific). Dithiothreitol (DTT, 10 mM final concentration) was added, and samples were incubated at 95°C with 800 rpm agitation for 10 min, followed by incubation with iodoacetamide (IAA, 55 mM final concentration) for 20 min in the dark. Samples were acetone precipitated (80% acetone final concentration) overnight at -20°C, then collected and washed twice in 80% acetone and resuspended in 8 M urea/40 mM HEPES for protein quantification using a bovine serum albumin (BSA) tryptophan assay (53). Samples were diluted in 50 mM ammonium bicarbonate and normalized to 50 μg protein for overnight LysC/trypsin digestion (Promega, protein:enzyme ratio, 50:1). Trifluoroacetic acid (TFA, 10% v/v) was added to quench the digestion, and peptides were purified using C18 Stop And Go Extraction (STAGE) tips (54).

Secretome sample preparation was performed using an in-solution digestion as previously described (52). Briefly, culture supernatant was filtered to remove whole cells and cellular debris by 0.22 μm syringe filters and incubated at 95°C for 10 min, followed by the addition of one-third volume of 8 M urea/40 mM HEPES to filtered sample. Samples were ultrasonicated in ice bath (15 cycles, 30 s on/30 s off) then reduced and alkylated with DTT and IAA, respectively. Samples were then enzymatically digested overnight, and STAGE-tip purified.

### Liquid chromatography-tandem mass spectrometry

Liquid chromatography-tandem mass spectrometry was performed as previously described with some modifications (15). Lyophilized peptides were resuspended in buffer A (0.1% formic acid) and analyzed on an Orbitrap Exploris 240 hybrid quadrupole-orbitrap mass spectrometry (Thermo Fisher Scientific) coupled to an Easy-nLC 1200 high-performance liquid chromatography device (Thermo Fisher Scientific). Resuspended samples were first loaded and separated on an in-line PepMap RSLC EASY-Spray column (75 μm by 50 cm) filled with C_18_ reverse-phase silica beads (2 μm) (Thermo Fisher Scientific). Peptides were subsequently electrosprayed into the mass spectrometer instrument across a linear gradient of 0-32% buffer B (80% acetonitrile, 0.5% acetic acid) over a 110 min gradient, followed by 95% buffer B wash for 5 min, and a 5 min hold with 4% buffer B, with a 250 nL/min flow rate. The mass spectrometer cycled between one full scan and MS/MS scans of the Top10 abundant peaks. Full scans (*m/z* 400 to 1600) were captured in the Orbitrap mass analyzer with a resolution of 60,000 at 200 *m/z*.

### Data processing

Data analysis of the mass spectrometry .RAW data files was completed using MaxQuant software (version 2.1.3) (55). The search was performed using the integrated Andromeda search engine against the reference *C. neoformans* var. *grubii* serotype A (strain H99/ATCC 208821) proteome (7429 sequences; downloaded on 14 July 2022) from Uniprot (56). The following parameters were included for data processing: trypsin enzyme specificity with maximum two missed cleavages; minimum peptide length of seven amino acids; fixed modifications – carbamidomethylation of cysteine, variable modifications – methionine oxidative and N-acetylation of proteins. Peptide spectral matches were filtered with a target-decoy approach at a false-discovery (FDR) of 1% with a minimum of two peptides required for protein identification. Relative label-free quantification (LFQ) and match between runs was enabled and the MaxLFQ algorithm used a minimum ratio count of one (57).

### Bioinformatics

Statistical analysis and data visualization was completed using Perseus (version 1.6.14) (58). Data was filtered for reverse database hits, contaminants, and proteins only identified by site. LFQ intensities were log_2_ transformed and filtered for valid values (three of four replicates in at least one group), followed by imputation of missing values from the normal distribution (width, 0.3; downshift, 1.8 standard deviations). Significant differences were evaluated by a Student’s *t* test (*p* ≤ 0.05) with multiple-hypothesis testing correction using the Benjamini-Hochberg FDR at 0.05 with S_0_ = 1 (59). A 1D annotation enrichment (i.e., tests for each annotation term whether the respective numerical values have a preference to be larger or smaller than the global distribution of the values for all proteins) was performed based on GOBP terms described with an FDR threshold of 0.05 using Benjamini-Hochberg method (23). Visualization of protein networks was performed using STRING analysis as described at https://string-db.org/.

### Disk Diffusion assay

The susceptibility of the *C. neoformans* strains to oxidative stress was performed as previously described (14,15). *C. neoformans* strains were grown to mid-log phase in YPD at 37°C; 2.5 × 10^5^ cells were plated with a cotton swab on semi-solid YPD. Sterile filter discs (Whatman MM, 10 mm diameter) were placed in plate center and 15 μL of 12% H_2_O_2_ was added to the disc. Plates were incubated at 37°C for 48 h, photographed, and measurements were taken from three locations to the nearest mm to determine the radius of the zone of inhibition. Experiment completed with four biological replicates and in technical duplicate.

### Cell culture

BALB/c WT immortalized macrophages (generously provided by Dr. Felix Meissner, Max Planck Institute of Biochemistry, Germany) were maintained at 37°C in 5% CO_2_ in Dulbecco’s Modified Eagle’s Medium (DMEM) supplemented with 10% heat-inactivated fetal bovine serum (FBS; Thermo Fisher), 2 mM Glutamax, 1% sodium pyruvate, 1% L-glutamine, and 5% pen/strep. For CFU assays, macrophages were seeded in 12-well plates at 0.1 × 10^6^ cells/well and grown until 70-80% confluence (i.e., 0.5 × 10^6^ cells/well). For fluorescence microscopy assays, 0.3 × 10^6^ cells were seeded in 6-well plates and grown until 70-80% confluence (i.e., 1.2 × 10^6^ cells/well).

### C. neoformans colony forming unit counts & fluconazole protection assays

A *C. neoformans* infection and fluconazole protection assay was performed as previously described with modifications (33). Briefly, *C. neoformans* strains were grown to mid-log phase in YPD at 30°C, collected, and washed twice in PBS. Macrophages were infected at a multiplicity of infection (MOI) of 5:1 (fungi:macrophage) in DMEM with pen/strep for 3 h at 37°C at 5% CO_2_. Following co-culture, infected cells were extensively washed six times with PBS to remove any adhered or non-phagocytosed fungal cells and fresh media supplemented with 20 μg/mL fluconazole was added for the remainder of the assay. At the indicated time points (i.e., 12 and 24 h), culture media containing extracellular fungal cells was collected and the infected macrophages were washed six times with PBS and lysed with 0.5% Tween-20 at room temperature for 10 min. The removed media containing extracellular fungal cells was centrifuged at 1200 × g for 12 min. The media containing fluconazole was removed and the cell pellet was resuspended in PBS. Serial dilutions of both resuspended extracellular fungal cells from culture media and lysed intracellular fungal cells were performed followed by plating on YPD and incubation for 48 h at 30°C.

To measure the number of fungal cells engulfed at the starting time point, 3 h post-inoculation (h.p.i.), the cell lysis as described above was performed. The 12 and 24 h.p.i collection time points consisted of both extracellular and intracellular CFU assessment, as described above. Experiment completed with three biological replicates and in technical duplicate.

To calculate fungal intracellular replication efficiency, for experiments that began at the initiation of infection, t_0_ (i.e., 3 h) and advance to the indicated time points, t_n_ (i.e., 24 h), the following formula was applied (33):

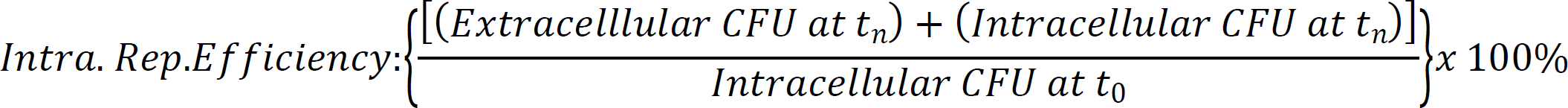

### Dilution plate assay

To analyze *C. neoformans* Wos2 response to heat, oxidative, osmotic and cell-wall stressors, dilution plate assays were performed as previously described (25,49). To assess osmotic stressor phenotypes, NaCl (1.5 M) and KCl (1.5 M) were supplemented in YPD medium. Cell wall and osmotic stressor phenotypes were assessed by adding caffeine (1 mg/mL), SDS (0.01%), or H_2_O_2_ (3 mM) to YPD medium. *C. neoformans* strains were grown to mid-log phase in YPD at 37°C and serial diluted in ten-fold (10^6^ cells/5 μL) on YPD plates supplemented with the different stressors and incubated at 37°C, unless otherwise stated. Images were taken every 24 h for 3 d. Experiment completed with two biological and technical replicates.

### Immunofluorescence

BALB/c WT immortalized macrophages were maintained as described above, to minimize autofluorescence, cells were grown in FluoroBrite DMEM media (Thermo Fisher Scientific) (supplemented with 10% FBS and 1% L-glutamine) 24 h prior to infection and throughout the infection protocol. Macrophages were infected at a MOI of 100 for 3 h at 37°C at 5% CO_2_, and samples were collected following co-culture and two washes with PBS. Protocol for immunostaining was adapted from previously described methods (60–62). Briefly, cells were fixed overnight in 4% paraformaldehyde in PBS at 4°C and plated onto 0.1% poly-L-lysine (Sigma-Aldrich) coated cell slide. Cells were incubated with buffer (0.1 M sodium citrate, 1.1 M sorbitol pH 5.5) containing 10 mg/mL of lytic enzymes (Sigma, L1412) and a protease inhibitor cocktail tablet and incubated at 30°C for 2 h. Slides were submerged in 99% methanol, followed by 100% acetone, and then blocked with 2% goat serum, 2% BSA, and 0.1% saponin in PBS for 1 h. Cells were incubated with monoclonal anti-FLAG M2 antibody (Sigma-Aldrich) diluted 1:100 in blocking solution for 1 h, followed by 1 h incubation with Anti-mouse Alexa Fluor 488 (AF488; Invitrogen) diluted to 1:200 containing DAPI (10 µg/mL). Slides were washed three times with PBS containing 0.01% Tween-20 between incubation steps. Coverslips were mounted with a drop of *SlowFade^TM^* Gold antifade (Life technologies).

Slides were imaged using a Leica DM5500B microscope, equipped with a Hamamatsu 3CCD digital camera operated through Volocity software ver. 6.3 (Quorum Technologies). A fixed exposure of 979ms was used to detect Alexa Fluor 488 bound to fungal cells, 49 ms used to detect DAPI, and 59 ms for phase contrast.

### Murine survival assay and tissue burden analysis

Murine infection assays were performed under approval of the Animal Utilization Protocol 4193 at the University of Guelph and accordance with all animal handling guidelines. *C. neoformans* strains (i.e., WT, *wos2*Δ) were grown overnight in YPD at 30°C, sub-cultured overnight at 1:100 in YPD, collected and washed in PBS twice, and resuspended at 4.0 × 10^6^ cells/mL in PBS. Ten female BALB/c elite mice aged 6-to-8-weeks old (Charles River Laboratories, ON, Canada) were intranasally inoculated with 50 μL of the *C. neoformans* cell suspension (inoculum of 2 × 10^5^ cells) under isoflurane anesthesia. The mice were monitored daily for signs of morbidity and euthanized by isoflurane and CO_2_ inhalation upon reaching endpoint specifications. Endpoint determining criteria includes loss of 20% total body weight, respiratory issues, or visible signs of neurological deficits. Tissue collection of lungs, bronchoalveolar lavage, and brain were retrieved upon termination. The collected tissues were weighed and homogenized in 1 mL PBS using a Bullet Blender Storm (Next Advance, Troy, NY, USA). Serial dilutions of the homogenized tissues were plated on YPD medium supplemented with 32 μg/mL chloramphenicol (provider) and incubated for 48 h at 30°C. All animal experiments were performed in accordance with the Canadian Council on Animal Care guidelines and approved the University of Guelph’s Animal Care Committee (Animal Utilization Protocol 4193).

## Author Contributions

J.G.-M. conceptualized the study. B.B. & J.G.-M. designed the study. B.B., A.S., S.P., S.K., N.C., & M.M. performed experiments. B.B.& J.G.-M. performed data analysis. B.B. & J.G.-M. designed and developed figures. B.B., & J.G.-M. wrote the manuscript. B.B. & J.G.-M. edited the manuscript. All authors have read and approved the submitted manuscript.

## Funding

In support of this project, B.B. is funded with a Canada Graduate Scholarship Doctoral-NSERC. B.B. is a fellow with the NSERC CREATE EvoFunPath program. J.G.-M. received funding from the University of Guelph, Canadian Foundation for Innovation (CFI-JELF #38798), Natural Sciences and Engineering Research Council of Canada (NSERC) Discovery Grant (RGPIN-2020-04838), and Canadian Institutes of Health Research (Project Grant).

## Acknowledgments

Thank you to members of the Geddes-McAlister lab for their informative and constructive feedback on project design and manuscript preparation. We thank Dr. Cezar Khursigara (University of Guelph) for their technical assistance and use of their microscopy equipment.

## Data Availability

The. RAW and affiliated files will be deposited into the publicly available PRIDE partner database for the ProteomeXchange consortium.

## Conflicts of Interest

The authors declare no conflicts of interest.

## Supporting information

**S1 Table. Significant different proteins identified in cellular proteome analysis of WT and *wos2*Δ strains grown in YPD and LIM conditions.** List of significantly different fungal proteins identified in the cellular proteome of *C. neoformans* WT and *wos2*Δ strains grown in enriched (YPD) and infection-mimicking (LIM) conditions. Statistical analysis (-log_10_ *P-*value reported): Student’s *t* test, *P* < 0.05; FDR = 0.05; S_0_ = 1. “Difference” represents the difference in normalized LFQ intensities (log_2_) for WT versus *wos2*Δ samples.

**S2 Table. Significant different proteins identified in secretome analysis of WT and wos2Δ strains grown in LIM conditions.** List of significantly different fungal proteins identified in the secretome of C. neoformans WT and wos2Δ strains grown in infection-mimicking (LIM) conditions. Statistical analysis (-log10 P-value reported): Student’s t test, P < 0.05; FDR = 0.05; S0 = 1. “Difference” represents the difference in normalized LFQ intensities (log2) for WT versus wos2Δ samples.

**S3 Table. Strains used in this study.**

## Notes

### Competing Interest Statement

The authors have declared no competing interest.

## References

1. Rajasingham R, Smith RM, Park BJ, Jarvis JN, Govender NP, Chiller TM, et al. Global burden of disease of HIV-associated cryptococcal meningitis: an updated analysis. Lancet Infect Dis. 2017;17(8):873–81.

2. Rajasingham R, Govender NP, Jordan A, Loyse A, Shroufi A, Denning DW, et al. The global burden of HIV-associated cryptococcal infection in adults in 2020: a modelling analysis. Lancet Infect Dis. 2022; 22(12):1748–1755

3. Ball, B, Woroszchuk, E, Sukumaran, A, Afaq, A, Carruthers-Lay, D, Muselius, B, West, H, Gee, L, Langille, M, Geddes-McAlister J. Proteome and secretome profiling of zinc availability in Cryptococcus neoformans, identifies Wos2 as a subtle influencer of fungal virulence determinants. BMC Microbiol. 2021; 21: 341

4. Muñoz MJ, Bejarano ER, Daga RR, Jimenez J. The identification of Wos2, a p23 homologue that interacts with Wee1 and Cdc2 in the mitotic control of fission yeasts. Genetics. 1999; 153(4): 1561–1572

5. Echtenkamp FJ, Zelin E, Oxelmark E, Woo JI, Andrews BJ, Garabedian M, et al. Global Functional Map of the p23 Molecular Chaperone Reveals an Extensive Cellular Network. Mol Cell. 2011 Jul;43(2):229–41.

6. Biebl MM, Lopez A, Rehn A, Freiburger L, Lawatscheck J, Blank B, et al. Structural elements in the flexible tail of the co-chaperone p23 coordinate client binding and progression of the Hsp90 chaperone cycle. Nat Commun. 2021 Dec 5;12(1):828.

7. Zhao R, Davey M, Hsu Y-C, Kaplanek P, Tong A, Parsons AB, et al. Navigating the Chaperone Network: An Integrative Map of Physical and Genetic Interactions Mediated by the Hsp90 Chaperone. Cell. 2005 Mar;120(5):715–27.

8. Horianopoulos LC, Kronstad JW. Chaperone Networks in Fungal Pathogens of Humans. J Fungi. 2021 Mar 12;7(3):209.

9. Sahasrabudhe P, Rohrberg J, Biebl MM, Rutz DA, Buchner J. The Plasticity of the Hsp90 Co-chaperone System. Mol Cell. 2017 Sep;67(6):947–961.e5.

10. Noddings CM, Wang RY-R, Johnson JL, Agard DA. Structure of Hsp90–p23–GR reveals the Hsp90 client-remodelling mechanism. Nature. 2022 Jan 20;601(7893):465–9.

11. Gu X, Xue W, Yin Y, Liu H, Li S, Sun X. The Hsp90 Co-chaperones Sti1, Aha1, and P23 Regulate Adaptive Responses to Antifungal Azoles. Front Microbiol. 2016 Oct 5;7.

12. Vartivarian SE, Anaissie EJ, Cowart RE, Sprigg HA, Tingler MJ., Jacobson ES. Regulation of Cryptococcal Capsular Polysaccharide by Iron. J Infect Dis. 1993 Jan 1;167(1):186–90.

13. Zaragoza O. Basic principles of the virulence of Cryptococcus. Virulence. 2019;10(1):490–501.

14. Giles SS, Stajich JE, Nichols C, Gerrald QD, Alspaugh JA, Dietrich F, et al. The Cryptococcus neoformans Catalase Gene Family and Its Role in Antioxidant Defense. Eukaryot Cell. 2006 Sep;5(9):1447–59.

15. Sukumaran A, Ball B, Krieger JR, Geddes-McAlister J. Cross-Kingdom Infection of Macrophages Reveals Pathogen- and Immune-Specific Global Reprogramming and Adaptation. Alspaugh JA, editor. MBio. 2022 Aug 30;13(4).

16. Idnurm A, Giles SS, Perfect JR, Heitman J. Peroxisome Function Regulates Growth on Glucose in the Basidiomycete Fungus Cryptococcus neoformans. Eukaryot Cell. 2007 Jan;6(1):60–72.

17. Galani K, Nissan TA, Petfalski E, Tollervey D, Hurt E. Rea1, a Dynein-related Nuclear AAA-ATPase, Is Involved in Late rRNA Processing and Nuclear Export of 60 S Subunits. J Biol Chem. 2004 Dec;279(53):55411–8.

18. Beach DL, Bloom K. ASH1 mRNA Localization in Three Acts. Koshland D, editor. Mol Biol Cell. 2001 Sep;12(9):2567–77.

19. Garbarino JE, Gibbons IR. Expression and genomic analysis of midasin, a novel and highly conserved AAA protein distantly related to dynein. BMC Genomics. 2002 Dec 8;3(1):18.

20. Stoldt S, Wenzel D, Hildenbeutel M, Wurm CA, Herrmann JM, Jakobs S. The inner-mitochondrial distribution of Oxa1 depends on the growth conditions and on the availability of substrates. Fox TD, editor. Mol Biol Cell. 2012 Jun 15;23(12):2292–301.

21. Szklarczyk D, Gable AL, Lyon D, Junge A, Wyder S, Huerta-Cepas J, et al. STRING v11: Protein-protein association networks with increased coverage, supporting functional discovery in genome-wide experimental datasets. Nucleic Acids Res. 2019; 47(D1): D607–613

22. Cadieux B, Lian T, Hu G, Wang J, Biondo C, Teti G, et al. The Mannoprotein Cig1 Supports Iron Acquisition From Heme and Virulence in the Pathogenic Fungus Cryptococcus neoformans. J Infect Dis. 2013 Apr 15;207(8):1339–47. A

23. Cox J, Mann M. 1D and 2D annotation enrichment: a statistical method integrating quantitative proteomics with complementary high-throughput data. BMC Bioinformatics. 2012; 13 (S12)

24. Xie JL, Bohovych I, Wong EOY, Lambert J-P, Gingras A-C, Khalimonchuk O, et al. Ydj1 governs fungal morphogenesis and stress response, and facilitates mitochondrial protein import via Mas1 and Mas2. Microb Cell. 2017 Oct 2;4(10):342–61.

25. Esher SK, Ost KS, Kohlbrenner MA, Pianalto KM, Telzrow CL, Campuzano A, et al. Defects in intracellular trafficking of fungal cell wall synthases lead to aberrant host immune recognition. May RC, editor. PLOS Pathog. 2018 Jun 4;14(6):e1007126.

26. Coelho C, Bocca AL, Casadevall A. The Intracellular Life of Cryptococcus neoformans. Annu Rev Pathol Mech Dis. 2014 Jan 24;9(1):219–38.

27. Cox GM, Harrison TS, McDade HC, Taborda CP, Heinrich G, Casadevall A, et al. Superoxide Dismutase Influences the Virulence of Cryptococcus neoformans by Affecting Growth within Macrophages. Infect Immun. 2003 Jan;71(1):173–80.

28. Rodrigues ML, Nakayasu ES, Oliveira DL, Nimrichter L, Nosanchuk JD, Almeida IC, et al. Extracellular vesicles produced by Cryptococcus neoformans contain protein components associated with virulence. Eukaryot Cell. 2008; 7(1): 58–67.

29. Reese AJ, Yoneda A, Breger JA, Beauvais A, Liu H, Griffith CL, et al. Loss of cell wall alpha(1-3) glucan affects Cryptococcus neoformans from ultrastructure to virulence. Mol Microbiol. 2007 Mar;63(5):1385–98.

30. Prole DL, Taylor CW. Identification and Analysis of Cation Channel Homologues in Human Pathogenic Fungi. Harris S, editor. PLoS One. 2012 Aug 2;7(8):e42404.

31. Stie J, Bruni G, Fox D. Surface-Associated Plasminogen Binding of Cryptococcus neoformans Promotes Extracellular Matrix Invasion. Lin X, editor. PLoS One. 2009 Jun 3;4(6):e5780.

32. Chatterjee S, Tatu U. Heat shock protein 90 localizes to the surface and augments virulence factors of Cryptococcus neoformans. Reynolds TB, editor. PLoS Negl Trop Dis. 2017 Aug 4;11(8):e0005836.

33. Qin Q-M, Luo J, Lin X, Pei J, Li L, Ficht TA, et al. Functional Analysis of Host Factors that Mediate the Intracellular Lifestyle of Cryptococcus neoformans. Andrianopoulos A, editor. PLoS Pathog. 2011 Jun 16;7(6):e1002078.

34. Fu C, Beattie SR, Jezewski AJ, Robbins N, Whitesell L, Krysan DJ, et al. Genetic analysis of Hsp90 function in Cryptococcus neoformans highlights key roles in stress tolerance and virulence. Stajich J, editor. Genetics. 2022 Jan 4;220(1).

35. Silveira CP, Piffer AC, Kmetzsch L, Fonseca FL, Soares DA, Staats CC, et al. The heat shock protein (Hsp) 70 of Cryptococcus neoformans is associated with the fungal cell surface and influences the interaction between yeast and host cells. Fungal Genet Biol. 2013 Nov;60:53–63.

36. Cowen LE, Singh SD, Köhler JR, Collins C, Zaas AK, Schell WA, et al. Harnessing Hsp90 function as a powerful, broadly effective therapeutic strategy for fungal infectious disease. Proc Natl Acad Sci. 2009 Feb 24;106(8):2818–23.

37. Zhang S, Hacham M, Panepinto J, Hu G, Shin S, Zhu X, et al. The Hsp70 member, Ssa1, acts as a DNA-binding transcriptional co-activator of laccase in Cryptococcus neoformans. Mol Microbiol. 2006 Nov;62(4):1090–101.

38. Eastman AJ, He X, Qiu Y, Davis MJ, Vedula P, Lyons DM, et al. Cryptococcal Heat Shock Protein 70 Homolog Ssa1 Contributes to Pulmonary Expansion of Cryptococcus neoformans during the Afferent Phase of the Immune Response by Promoting Macrophage M2 Polarization. J Immunol. 2015 Jun 15;194(12):5999–6010.

39. Cowen LE, Carpenter AE, Matangkasombut O, Fink GR, Lindquist S. Genetic architecture of Hsp90-dependent drug resistance. Eukaryot Cell. 2006 Dec;5(12):2184–8.

40. Cowen LE, Lindquist S. Hsp90 Potentiates the Rapid Evolution of New Traits: Drug Resistance in Diverse Fungi. Science. 2005 Sep 30;309(5744):2185–9.

41. Shapiro RS, Uppuluri P, Zaas AK, Collins C, Senn H, Perfect JR, et al. Hsp90 orchestrates temperature-dependent Candida albicans morphogenesis via Ras1-PKA signaling. Curr Biol. 2009 Apr 28;19(8):621–9.

42. Cowen LE. The fungal Achilles’ heel: targeting Hsp90 to cripple fungal pathogens. Curr Opin Microbiol. 2013 Aug;16(4):377–84.

43. Cordeiro R de A, Evangelista AJ de J, Serpa R, Marques FJ de F, Melo CVS de, Oliveira JS de, et al. Inhibition of heat-shock protein 90 enhances the susceptibility to antifungals and reduces the virulence of Cryptococcus neoformans/Cryptococcus gattii species complex. Microbiology. 2016 Feb 1;162(2):309–17.

44. Forafonov F, Toogun OA, Grad I, Suslova E, Freeman BC, Picard D. p23/Sba1p Protects against Hsp90 Inhibitors Independently of Its Intrinsic Chaperone Activity. Mol Cell Biol. 2008 May 15;28(10):3446–56.

45. Morano KA, Grant CM, Moye-Rowley WS. The Response to Heat Shock and Oxidative Stress in Saccharomyces cerevisiae. Genetics. 2012 Apr 1;190(4):1157–95.

46. Brown AJ, Haynes K, Quinn J. Nitrosative and oxidative stress responses in fungal pathogenicity. Curr Opin Microbiol. 2009 Aug;12(4):384–91.

47. Zaragoza O, Chrisman CJ, Castelli MV, Frases S, Cuenca-Estrella M, Rodríguez-Tudela JL, et al. Capsule enlargement in Cryptococcus neoformans confers resistance to oxidative stress suggesting a mechanism for intracellular survival. Cell Microbiol. 2008 Oct;10(10):2043–57.

48. Wang Y, Aisen P, Casadevall A. Cryptococcus neoformans melanin and virulence: Mechanism of action. Infect Immun. 1995;63(8):3131–6.

49. Leipheimer J, Bloom ALM, Campomizzi CS, Salei Y, Panepinto JC. Translational Regulation Promotes Oxidative Stress Resistance in the Human Fungal Pathogen Cryptococcus neoformans. Alspaugh JA, editor. MBio. 2019 Dec 24;10(6).

50. Santiago-tirado FH, Onken MD, Cooper JA, Klein RS, Doering TL. Trojan Horse Transit Contributes to Blood-Brain Barrier Crossing of a Eukaryotic Pathogen. MBio. 2017; 8(1): e02183–16

51. Casadevall A, Coelho C, Alanio A. Mechanisms of Cryptococcus neoformans-mediated host damage. Front Immunol. 2018;9:1–8.

52. Ball B, Geddes-McAlister J. Quantitative Proteomic Profiling of Cryptococcus neoformans. Curr Protoc Microbiol. 2019;55(1):1–15.

53. Wiśniewski JR, Gaugaz FZ. Fast and Sensitive Total Protein and Peptide Assays for Proteomic Analysis. Anal Chem. 2015 Apr 21;87(8):4110–6.

54. Rappsilber J, Mann M, Ishihama Y. Protocol for micro-purification, enrichment, pre-fractionation and storage of peptides for proteomics using StageTips. Nat Protoc. 2007 Aug 2;2(8):1896–906.

55. Cox J, Mann M. MaxQuant enables high peptide identification rates, individualized p.p.b.-range mass accuracies and proteome-wide protein quantification. Nat Biotechnol. 2008 Dec 30;26(12):1367–72.

56. Cox J, Neuhauser N, Michalski A, Scheltema RA, Olsen J V., Mann M. Andromeda: A Peptide Search Engine Integrated into the MaxQuant Environment. J Proteome Res. 2011 Apr;10(4):1794–805.

57. Cox J, Hein MY, Luber CA, Paron I, Nagaraj N, Mann M. Accurate Proteome-wide Label-free Quantification by Delayed Normalization and Maximal Peptide Ratio Extraction, Termed MaxLFQ. Mol Cell Proteomics. 2014 Sep;13(9):2513–26.

58. Tyanova S, Temu T, Sinitcyn P, Carlson A, Hein MY, Geiger T, et al. The Perseus computational platform for comprehensive analysis of (prote)omics data. Nat Methods. 2016 Sep 27;13(9):731–40.

59. Benjamini Y, Hochberg Y. Controlling the False Discovery Rate: A Practical and Powerful Approach to Multiple Testing. J R Stat Soc Ser B. 1995; 43(5): 2055–2085

60. Garcia-Santamarina S, Probst C, Festa RA, Ding C, Smith AD, Conklin SE, et al. A lytic polysaccharide monooxygenase-like protein functions in fungal copper import and meningitis. Nat Chem Biol. 2020;16(3):337–44.

61. Hiraga S, Ichinose C, Niki H, Yamazoe M. Cell Cycle–Dependent Duplication and Bidirectional Migration of SeqA-Associated DNA–Protein Complexes in. Mol Cell. 1998 Feb;1(3):381–7.

62. Müller P, Chikkaballi D, Hensel M. Functional Dissection of SseF, a Membrane-Integral Effector Protein of Intracellular Salmonella enterica. Webber MA, editor. PLoS OneS. 2012 Apr 18;7(4):e35004.

